# Behavioral and Pharmacological Validation of the Differential Reinforcement of Low-Rate Behavior Paradigm in Non-Human Primates

**DOI:** 10.1101/2025.06.02.657510

**Authors:** Casey R. Vanderlip, Shelby R. Dunn, Jennifer L. Basile, Joseph G. Wettstein, Jeffrey A. Vivian, Courtney Glavis-Bloom

## Abstract

Depression remains a leading cause of disability worldwide, yet the predictive validity of many preclinical behavioral assays for antidepressant efficacy remains limited. The Differential Reinforcement of Low-Rate Behavior (DRL) task has classically been used in rodents to identify antidepressant-like effects, but its utility in non-human primates (NHPs) has not been established. Here, we adapted the DRL task for use in adult male cynomolgus macaques (*Macaca fascicularis*) and evaluated its pharmacological sensitivity and translational relevance across 19 compounds spanning multiple drug classes. Antidepressants, including SSRIs, SNRIs, NRIs, NDRIs, TCAs, MAOIs, and PDE4 inhibitors, generally shifted DRL performance in an antidepressant-like direction, increasing reinforcers earned and inter-response times while decreasing response output. In contrast, benzodiazepine and antipsychotic control compounds did not produce a consistent antidepressant-like profile, whereas stimulant effects were mixed, with nicotine and cocaine also producing overlapping antidepressant-like behavioral effects. Importantly, the primate DRL task identified antidepressant-like effects of PDE4 inhibitors while also capturing emesis, a dose-limiting side effect not observable in rodent models. These findings support the primate DRL task as a translationally relevant platform for screening antidepressant-like efficacy, while also highlighting important design considerations for interpreting pharmacological sensitivity in the NHP setting. By modeling behavioral processes implicated in depression, including response inhibition and temporal regulation, this assay offers a unique opportunity to bridge preclinical and clinical antidepressant development with improved sensitivity to both efficacy and tolerability.

## 1. INTRODUCTION

Depression remains a leading cause of disability worldwide, yet translating promising preclinical findings into effective clinical therapies is still challenging (Willner, 1984; Wang et al., 2017; Lee et al., 2023; Moncrieff et al., 2023). While many benefit from current antidepressants, a significant subset remains treatment-resistant. Clinical failures of new compounds often stem from limited efficacy, poor tolerability, or both, reflecting a key bottleneck: lack of behavioral assays with strong predictive validity and translational relevance for novel therapeutic screening.

The Differential Reinforcement of Low-Rate Behavior (DRL) task is a classic preclinical assay used in rodents for identifying antidepressant-like effects of therapeutics (McGuire and Seiden, 1980; O’Donnell and Seiden, 1983, 1984; Seiden et al., 1985; Richards and Seiden, 1991; O’Donnell et al., 2005). In the DRL task, subjects are rewarded for withholding responses for a minimum, un-signaled delay interval. Premature responses reset the interval and forfeit reward. Successful performance therefore depends on coordinated response inhibition and temporal regulation, requiring subjects to withhold premature responses while estimating when sufficient time has elapsed to emit a reinforced response. As a result, improved DRL performance is reflected by increased reinforcers earned, reduced response output, and lengthened inter-response intervals. The combination of precise timing, response inhibition, and sustained attention required for optimal performance is often impaired in depression (Fales et al., 2009; Ma, 2015; Hsu et al., 2019). These behaviors are subserved by a distributed neural circuit involving the prefrontal cortex, striatum, and hippocampus, brain regions consistently implicated in the pathophysiology of depression (Cho and Jeantet, 2010; Liu et al., 2017; Hamilton et al., 2018; Belleau et al., 2019; Liao and Pattij, 2022). In addition, performance on the DRL task is modulated by serotonergic signaling, aligning it with the pharmacological targets of many clinically effective antidepressants (Liao and Pattij, 2022).

In rodents, the DRL paradigm has demonstrated strong predictive validity for antidepressant efficacy (Marek and Salek, 2020; Marek et al., 2016). Compounds across several pharmacological classes, including selective serotonin reuptake inhibitors (SSRIs), tricyclic antidepressants (TCAs), monoamine oxidase inhibitors (MAOIs), consistently improve DRL performance. These improvements are reflected by increased number of reinforcers obtained, decreased total responding, and increased lengths of inter-response intervals compared to control animals (McGuire and Seiden, 1980; O’Donnell and Seiden, 1983; Richards and Seiden, 1991; O’Donnell et al., 2005). In contrast, other psychotropics such as anxiolytics, sedatives, and stimulants do not enhance performance, supporting the utility of DRL for antidepressant screening (Marek et al., 1993; O’Donnell and Seiden, 1982).

Despite this, the DRL paradigm has not yet been validated in non-human primates (NHPs). This is a significant gap, given that rodents differ significantly from humans in terms of genetics, neuroanatomy, pharmacokinetics, and behavioral complexity. NHPs share closer homology in cortical organization and neurotransmitter systems (Nestler et al., 2002; Liu et al., 2017; Hamilton et al., 2018). Importantly, NHPs exhibit depressive-like phenotypes under naturalistic conditions, enhancing their translational relevance (Xu et al., 2015; Deng et al., 2019; Price et al., 1994). Moreover, NHPs display clinically relevant side effects, including emesis, which rodents cannot. This is a significant limitation when evaluating the tolerability of drug candidates such as phosphodiesterase-4 (PDE4) inhibitors (Robichaud et al., 2001; McDonough et al., 2020).

Here, we adapted the DRL paradigm for macaque monkeys and conducted a comprehensive pharmacological validation of the model. The primary objectives were to (1) evaluate the sensitivity of NHP DRL performance to established antidepressants, (2) assess the specificity of the task using psychotropic control compounds, and (3) determine whether the model could detect emesis and other side effects relevant to clinical translation. We tested 19 pharmacotherapies from multiple classes, including SSRIs, serotonin-norepinephrine reuptake inhibitors (SNRIs), noradrenergic reuptake inhibitors (NRIs), a norepinephrine-dopamine reuptake inhibitor (NDRI), a TCA, a MAOI, a benzodiazepine (BZD), an atypical antipsychotic, several psychostimulants, and several phosphodiesterase-4 (PDE4) inhibitors for activity in the DRL task. This allowed us to evaluate the translational validity of the NHP DRL model and to examine how well the task distinguishes antidepressant-like effects from those of other psychoactive compounds. By establishing the validity, sensitivity, and translational utility of the DRL paradigm in macaques, this study provides a foundation for its use in preclinical evaluation of novel antidepressant therapeutics, particularly for mechanistically distinct or side-effect-prone drug classes.

## 2. MATERIALS AND METHODS

### 2.1. Subjects

Eight adult male cynomolgus macaques (*Macaca fascicularis*) weighing 6–10 kg were used in this study. The exact sample size for each of the compounds tested is reported throughout the Results section. Sample sizes were determined a priori based on typical group sizes for non-human primate pharmacology studies. Animals were housed in a same-sex colony room maintained on a 12-hour light/dark cycle and at 21 ± 2 **°**C, 40 ± 10% humidity. Monkeys were pair-housed except for during drug administration, feeding, nights, and weekends. Water was available *ad libitum*, and animals were fed a standard diet (Purina High Protein #5045) at the end of each day. Environmental enrichment, including toys and fresh fruit and vegetables, was provided. Animals were in good health as determined by routine veterinary oversight. Prior to the present study, animals had been trained on operant procedures and had a prior history of drug exposures for other pharmacology investigations. All procedures were approved by the Institutional Animal Care and Use Committee at Roche Palo Alto and SRI International and adhered to the USDA Guide for the Care and Use of Laboratory Animals.

### 2.2. Apparatus

Behavioral testing was conducted in a sound-attenuated chamber (30”W, 30”D, 59”H; Med Associates, St. Albans, VT), equipped with a light, exhaust fan, and a white-noise generator. Subjects were seated in a standard Plexiglas restraint chair. A panel inside the chamber contained a retractable lever and indicator light that were activated at the beginning of the session and inactivated and retracted at the end. Reinforcers (300 mg banana-flavored pellet; TestDiet, Dean’s Animal Feed, Redwood City, CA) were delivered to the lower right of the chair from a feeder mounted on top of the chamber. The task and data acquisition were controlled using Graphic State software (Coulbourn Instruments, Allentown, PA).

### 2.3. Training and Testing Procedures

Monkeys were initially trained to press a lever for food reinforcement during 60-minute sessions conducted 4–5 times per week. Once reliable lever pressing was established under a fixed-ratio 1 (FR1) schedule where each lever press delivered one pellet, the animals were transitioned to a Differential Reinforcement of Low-Rate (DRL) schedule. During DRL testing, sessions were conducted twice weekly. Under this schedule, a response was reinforced only if it occurred after a minimum un-signaled delay had elapsed since the preceding response. Premature responses reset the delay interval and were not reinforced.

Because baseline response tendencies differed across subjects, the DRL delay was individualized during training to establish stable and measurable performance within each monkey. Animals were initially started on a short delay, and the delay was then progressively increased in a stepwise manner until stable performance was achieved, defined as earning 2–6 reinforcers for 3 consecutive sessions. The delay that supported this level of performance was then used for testing. Final assigned delays ranged from 80 to 240 seconds across subjects and drug studies. Prior to initiation of each drug study, baseline performance was re-evaluated and the delay was adjusted as needed to re-establish this criterion. This approach was used to account for the potential drift in performance over the course of the multi-year study, including changes related to experience and practice. Once established for a given drug study, each monkey’s DRL delay remained constant throughout testing for that compound, although the delay used could differ across monkeys and across studies conducted at different times.

### 2.4. Drugs

Drug doses and pretreatment times were selected based on available literature and internal pharmacokinetic data to target clinically relevant exposure levels. Paroxetine (0.3, 1, 3 mg/kg), fluoxetine (1, 3, 10 mg/kg), duloxetine (0.03, 0.1, 0.3, 1 mg/kg), atomoxetine (1, 3 mg/kg), desipramine (1, 3, 10 mg/kg), reboxetine (1, 3, 10 mg/kg), and bupropion (0.3, 1, 3 mg/kg) were all dissolved in water. Sibutramine (0.1, 0.18, 0.3, 0.56 mg/kg), moclobemide (0.3, 1, 3 mg/kg), chlordiazepoxide (1, 3, 10, 30 mg/kg), nicotine (0.03, 0.1, 0.18, 0.3 mg/kg), cocaine (0.03, 0.1, 0.3 mg/kg), methylphenidate (0.1, 0.18 mg/kg), and modafinil (0.1, 0.3, 1 mg/kg) were dissolved in saline. Citalopram (1, 3, 10 mg/kg) was dissolved in fruit punch. Olanzapine (0.1, 0.3 mg/kg) was dissolved in 0.3% Tween 80. Roflumilast (0.1, 0.3, 1 mg/kg), rolipram (0.01, 0.03, 0.1, 0.3 mg/kg), and cilomilast (1, 3 mg/kg) were dissolved in 10% Cremophor. All drugs were administered intramuscularly (i.m.) with an injection volume of 0.1 ml/kg, with the exception of citalopram and atomoxetine, which were dosed orally (p.o.) at a dose volume of 2.5 ml/kg. Pretreatment times were 1 hour except for citalopram at 4 hours, duloxetine at 5 hours, reboxetine, methylphenidate, roflumilast, rolipram, and cilomilast at 30 minutes, and olanzapine and nicotine at 15 minutes.

### 2.5. Experimental Design

Drug effects were evaluated in a repeated-measures within-subject design. Subjects were tested twice weekly, in 60-minute sessions, with the first weekly session always serving as a vehicle control session and the second weekly session involving drug or vehicle administration. Dose order within each compound was assigned using a manually-generated counterbalanced Latin square schedule prepared before the study began, such that each subject eventually received every dose while dose conditions were staggered across animals rather than administered to all subjects on the same day. The investigators performing the behavioral testing and data extraction were blinded to the treatment condition of the animals.

Washout between doses of the same compound was 1 week. Washout between compounds was at least 1 week. This design minimized acute carryover between sessions while allowing repeated within-subject assessment across compounds.

### 2.6. Data Analysis

Behavioral performance on the DRL task was evaluated using three session-level dependent variables: number of reinforcers obtained (RF), number of responses (RR), and inter-response time (IRT). For RF and RR, total session counts were calculated. IRT was defined as the average elapsed time, in seconds, between consecutive lever presses within a session. Specifically, the time between each pair of subsequent lever presses was calculated and averaged to generate one session-level IRT value per subject. The experimental unit for all analyses was the individual monkey. Investigators performing data analysis were not blinded to the experimental conditions.

A priori exclusion criteria for analysis were limited to sessions with fewer than six total responses. Sessions with extremely low response counts were considered insufficient to yield stable and interpretable estimates of DRL performance, particularly because the expected antidepressant-like profile includes reduced responding. Below this threshold, further decreases in responding approach floor, making it difficult to distinguish improved schedule control from nonspecific behavioral suppression or disengagement. This resulted in exclusion of a total of 41 sessions out of a total of 518 across all drugs, doses, and animals. Table 1 provides detailed exclusion information for each of the compounds and doses.

**Table 1.**
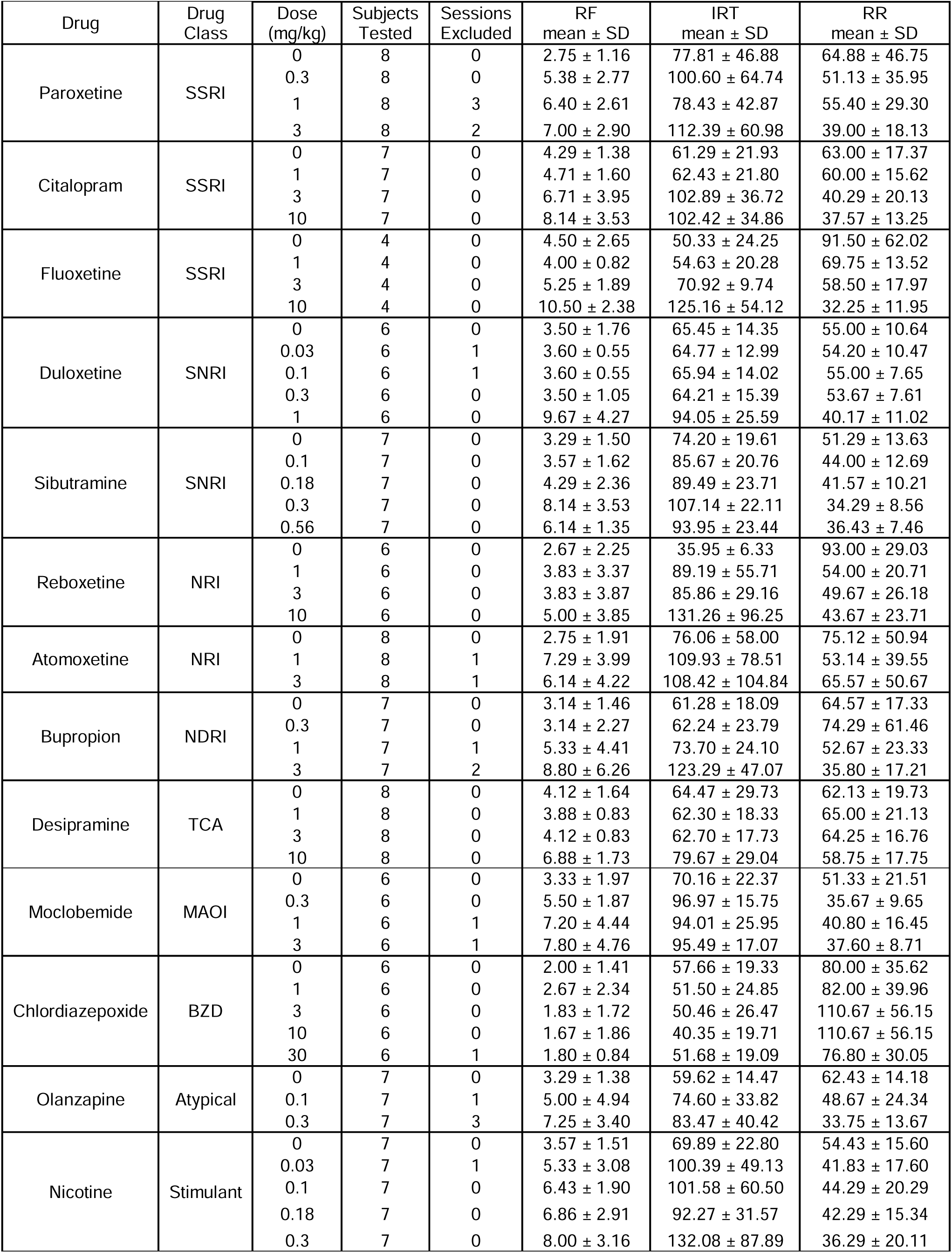
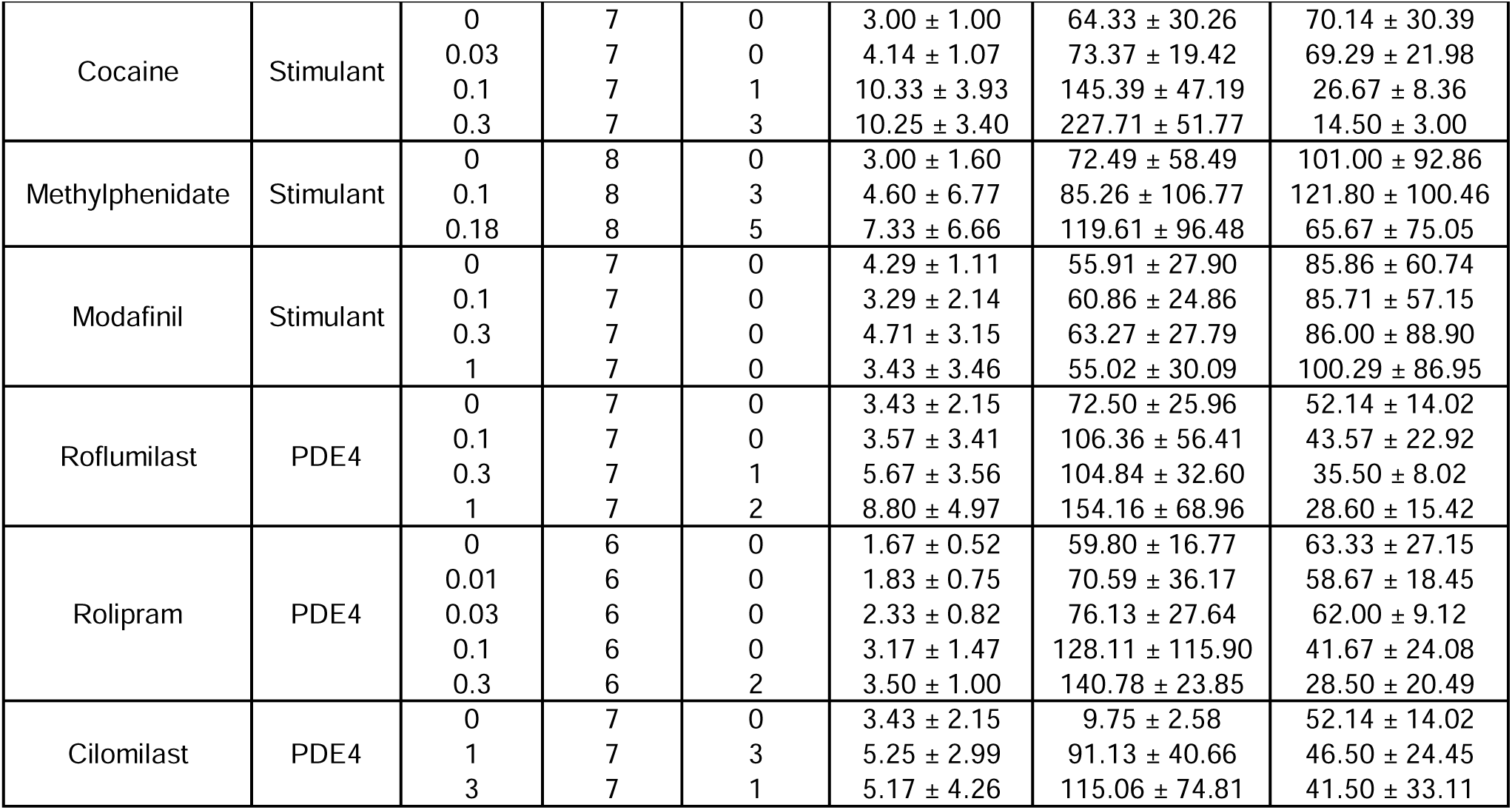
Group-level statistical summary. Abbreviations: SSRI - Selective serotonin reuptake inhibitor; SNRI - serotonin-norepinephrine reuptake inhibitor; NRI - norepinephrine reuptake inhibitor; NDRI - norepinephrine-dopamine reuptake inhibitor; TCA - tricyclic antidepressant; MAOI - monoamine oxidase inhibitor; BZD - benzodiazepine; PDE4 - phosphodiesterase-4 inhibitor; RF - reinforcers earned; RR - responses; IRT - inter-response time.

To account for repeated measures within subjects, non-normal session-level outcomes, and small sample sizes, we conducted rank-based linear mixed effects models for each drug and dependent variable. No additional parametric assumption testing was required beyond model specification. Models were fitted using the lmer() function in the lmerTest package in R (version 4.5.0), with Dose as a fixed effect and Monkey as a random intercept. Significance of the Dose effect was assessed via F-tests using Satterthwaite’s approximation for degrees of freedom. For pairwise post-hoc comparisons between doses, we used estimated marginal means and Tukey-adjusted contrasts via the emmeans package. Results are reported both for overall model significance (main effect of Dose) and for pairwise contrasts between doses and vehicle conditions. Significance was defined by α = 0.05. Group-level means for each compound and dose are reported in Table 1, and a summary of overall findings is reported in Table 2.

**Table 2.**
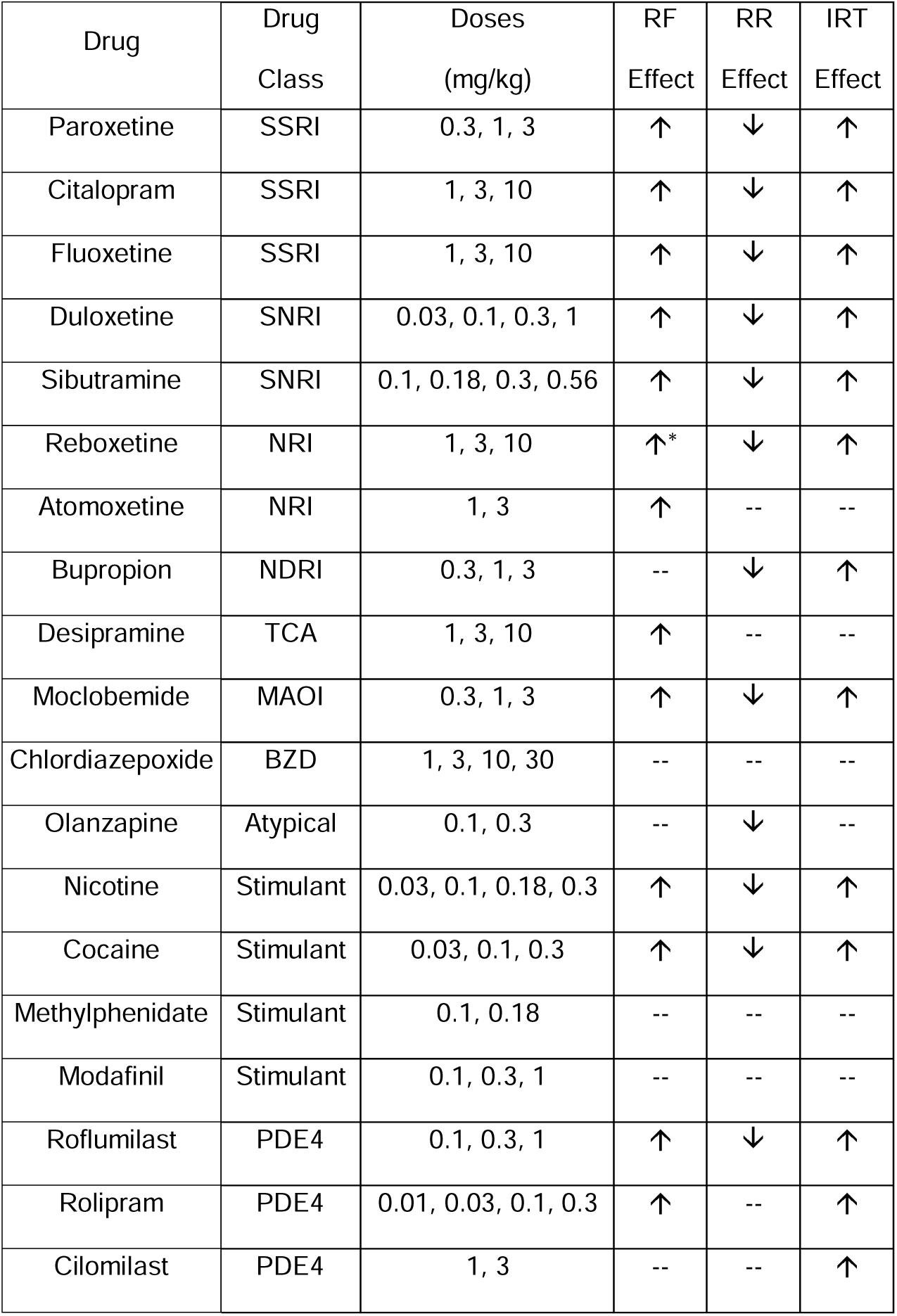
Drug dose ranges and summary of behavioral effects. Abbreviations: SSRI - Selective serotonin reuptake inhibitor; SNRI - serotonin-norepinephrine reuptake inhibitor; NRI - norepinephrine reuptake inhibitor; NDRI - norepinephrine-dopamine reuptake inhibitor; TCA - tricyclic antidepressant; MAOI - monoamine oxidase inhibitor; BZD - benzodiazepine; PDE4 - phosphodiesterase-4 inhibitor; IM - intramuscular; PO - per oral; RF - reinforcers earned; RR - responses; IRT - inter-response time. *approached significance (p = 0.06)

## 3. RESULTS

### 3.1. Pharmacological Validation of the NHP-DRL Assay

To evaluate the sensitivity of the primate DRL model to antidepressant compounds, we first examined drugs across mechanistically diverse antidepressant classes. These included SSRIs, SNRIs, selective norepinephrine reuptake inhibitors, norepinephrine-dopamine reuptake inhibitors, TCAs, and MAOIs.

#### 3.1.1. Selective Serotonin Reuptake Inhibitors (SSRIs)

Paroxetine (0.3–3 mg/kg; n = 8) produced a dose-dependent increase in RF (Figure 1a; F(3, 17.6) = 7.11, p = 0.002) and IRT (F(3, 16.1) = 7.22, p = 0.003), along with a significant decrease in RR (F(3, 16.1) = 4.49, p = 0.018). Post-hoc analyses revealed that each of the doses administered (0.3, 1, and 3 mg/kg) significantly increased RF, the two highest doses increased IRT, and the highest dose decreased RR (all p < 0.05). Similarly, citalopram (1–10 mg/kg; n = 7) significantly increased RF (Figure 1b; F(3, 18) = 3.34, p = 0.042) and IRT (F(3, 18) = 10.01, p < 0.001), and decreased RR (F(3, 18) = 8.22, p = 0.001). These effects were dose-dependent, with RF significantly increased at 10 mg/kg, whereas RR was significantly decreased and IRT significantly increased at both 3 and 10 mg/kg (all p < 0.05). Fluoxetine (1–10 mg/kg; n = 4) also produced significant increases in RF (Figure 1c; F(3, 9) = 9.62, p = 0.004) and IRT (F(3, 9) = 13.5, p = 0.001), and a significant decrease in RR (F(3, 9) = 11.59, p = 0.002). Post-hoc analyses revealed that the highest dose (10 mg/kg) significantly altered performance compared with vehicle administration for all three dependent variables (all p < 0.05).

**Figure 1.**
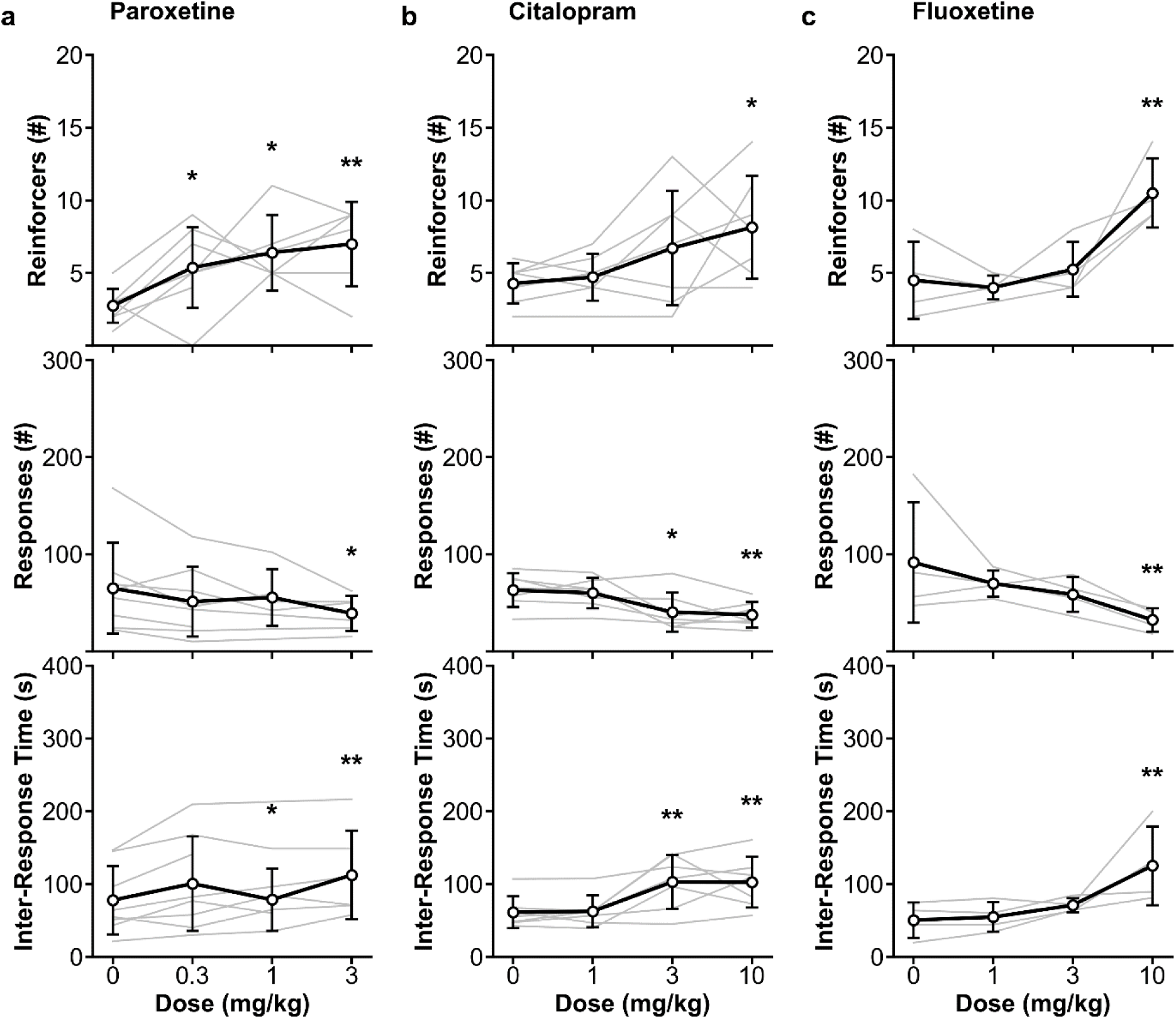
Effects of SSRIs on DRL task performance. Reinforcers earned, number of responses, and inter-response time following acute administration of (a) paroxetine, (b) citalopram, and (c) fluoxetine. All three SSRIs shifted DRL performance in an antidepressant-like direction, with increases in reinforcers and inter-response time and decreases in number of responses. Black lines are group means ± SD. Grey lines are individual animals. *p < 0.05 vs vehicle. **p < 0.01 vs vehicle.

#### 3.1.2. SNRI, NRI, NDRI, TCA, and MAOI Antidepressants

We tested the effects of two serotonin-norepinephrine reuptake inhibitors (SNRIs): duloxetine and sibutramine. Duloxetine (0.03–1 mg/kg; n = 6) administration resulted in a marked increase in RF (Figure 2a; F(4, 17.6) = 8.12, p < 0.001) and IRT (F(4, 18.1) = 7.79, p < 0.001), and a decrease in RR (F(4, 18.1) = 5.17, p = 0.006). These effects were dose-dependent, with the 1.0 mg/kg dose reaching significance for each of the dependent variables (all p < 0.05). Administration of the SNRI sibutramine (0.1–0.56 mg/kg, n = 7) also significantly affected each of the dependent variables. There was a significant increase in RF (Figure 2b; F(4, 24) = 5.92, p = 0.002) and IRT (F(4, 24) = 3.24, p = 0.029), and a significant decrease in RR (F(4, 24) = 4.26, p = 0.010). Post-hoc analyses revealed that the 0.3 and 0.56 mg/kg doses significantly increased RF compared with vehicle administration, and that the 0.3 mg/kg dose significantly altered RR and IRT (all p < 0.05).

**Figure 2.**
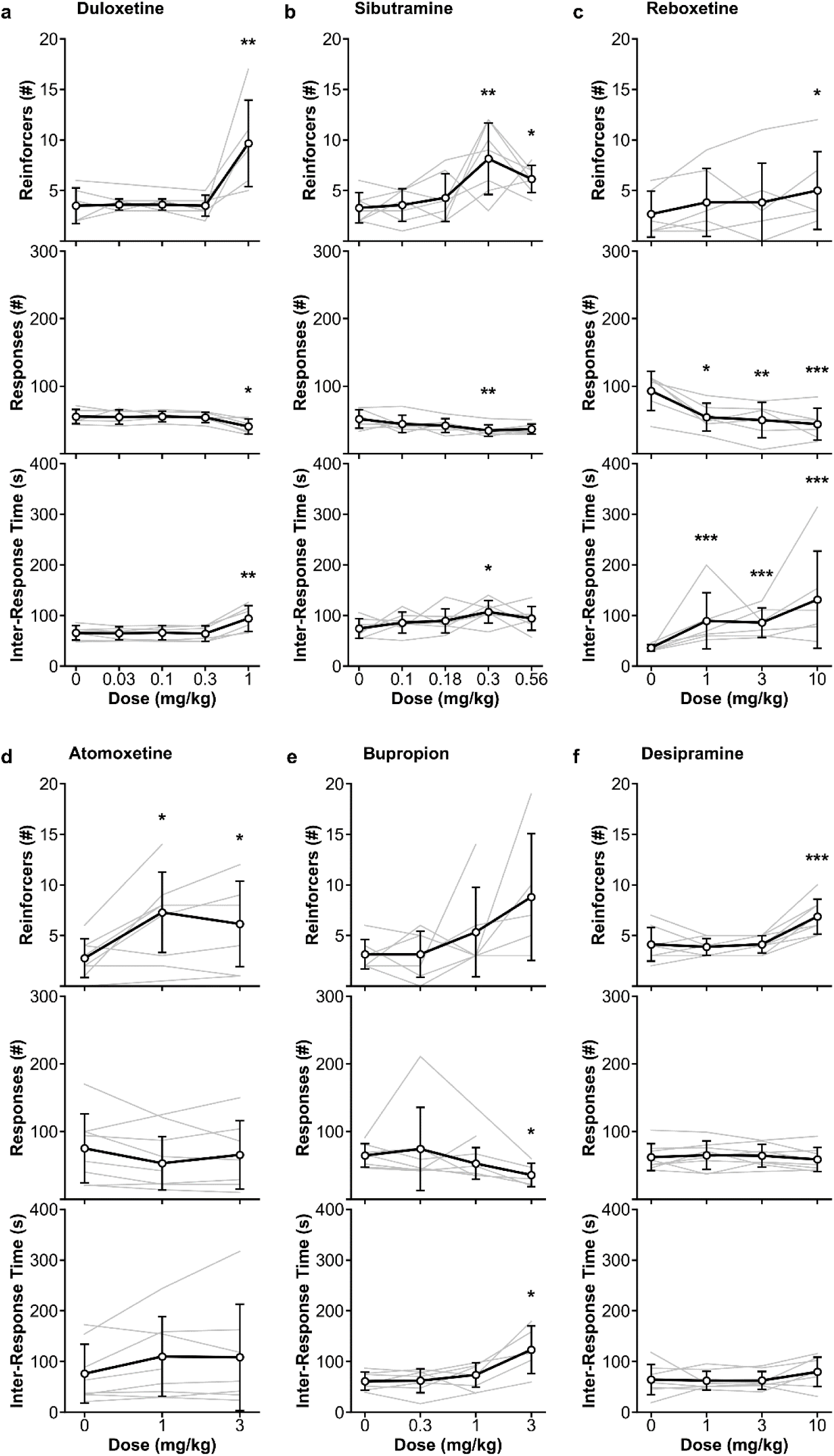
Behavioral effects of non-SSRI antidepressants in the primate DRL task. Reinforcers earned, number of responses, and inter-response time following acute administration of (a) duloxetine, (b) sibutramine, (c) reboxetine, (d) atomoxetine, (e) bupropion, and (f) desipramine. Black lines are group means ± SD. Grey lines are individual animals. *p < 0.05 vs vehicle. **p < 0.01 vs vehicle. ***p < 0.001 vs vehicle.

We tested the effects of two selective norepinephrine reuptake inhibitors (NRIs): reboxetine and atomoxetine. Reboxetine (1–10 mg/kg; n = 6) produced an effect on RF that approached significance (Figure 2c; F(3, 15) = 3.07, p = 0.060), and significant effects on both IRT (F(3, 15) = 23.69, p < 0.001) and RR (F(3, 15) = 10.30, p < 0.001). For RF, there was a significant effect of the 10 mg/kg dose compared to vehicle performance (p < 0.05). All three doses (1, 3, 10 mg/kg) significantly increased IRT and decreased RR compared to vehicle (all p < 0.05). Acute atomoxetine treatment (1–3 mg/kg; n = 8) had an overall significant effect on RF (Figure 2d; F(2, 11.6) = 5.54, p = 0.020), but no significant main effects on either IRT (F(2, 12.1) = 1.00, p = 0.397) or RR (F(2, 12.1) = 0.48, p = 0.628). Both the 1 and 3 mg/kg doses significantly increased RF compared to vehicle (p < 0.05).

The norepinephrine-dopamine reuptake inhibitor (NDRI), bupropion (0.3–3 mg/kg, n = 7) significantly increased IRT (F(3, 15.2) = 4.77, p = 0.016) and decreased RR (F(3, 15.3) = 3.49, p = 0.041), but did not significantly alter RF (Figure 2e; F(3, 21) = 2.61, p = 0.078). Post-hoc analyses revealed that 3 mg/kg bupropion significantly increased IRT relative to vehicle and significantly decreased RR at the same dose (all p < 0.05).

The tricyclic antidepressant (TCA), desipramine (1–10 mg/kg; n = 8) significantly increased RF (Figure 2f; F(3, 21) = 18.43, p < 0.001) but did not significantly affect IRT (F(3, 21) = 2.43, p = 0.094) or RR (F(3, 21) = 0.84, p = 0.489). The significant overall effects of desipramine on RF were driven by significant increases in performance at the 10 mg/kg dose (p < 0.05).

The monoamine oxidase inhibitor, moclobemide (0.3–3 mg/kg; n = 6) produced significant increases in both RF (Figure 3a; F(3, 12.7) = 4.52, p = 0.023) and IRT (F(3, 13.5) = 4.82, p = 0.017), and a significant decrease in RR (F(3, 13.3) = 4.64, p = 0.020). Significant increases in RF and decreases in RR were observed specifically at the 1 and 3 mg/kg doses (all p < 0.05). Significant increases in IRT were observed at all three doses tested (all p < 0.05).

**Figure 3.**
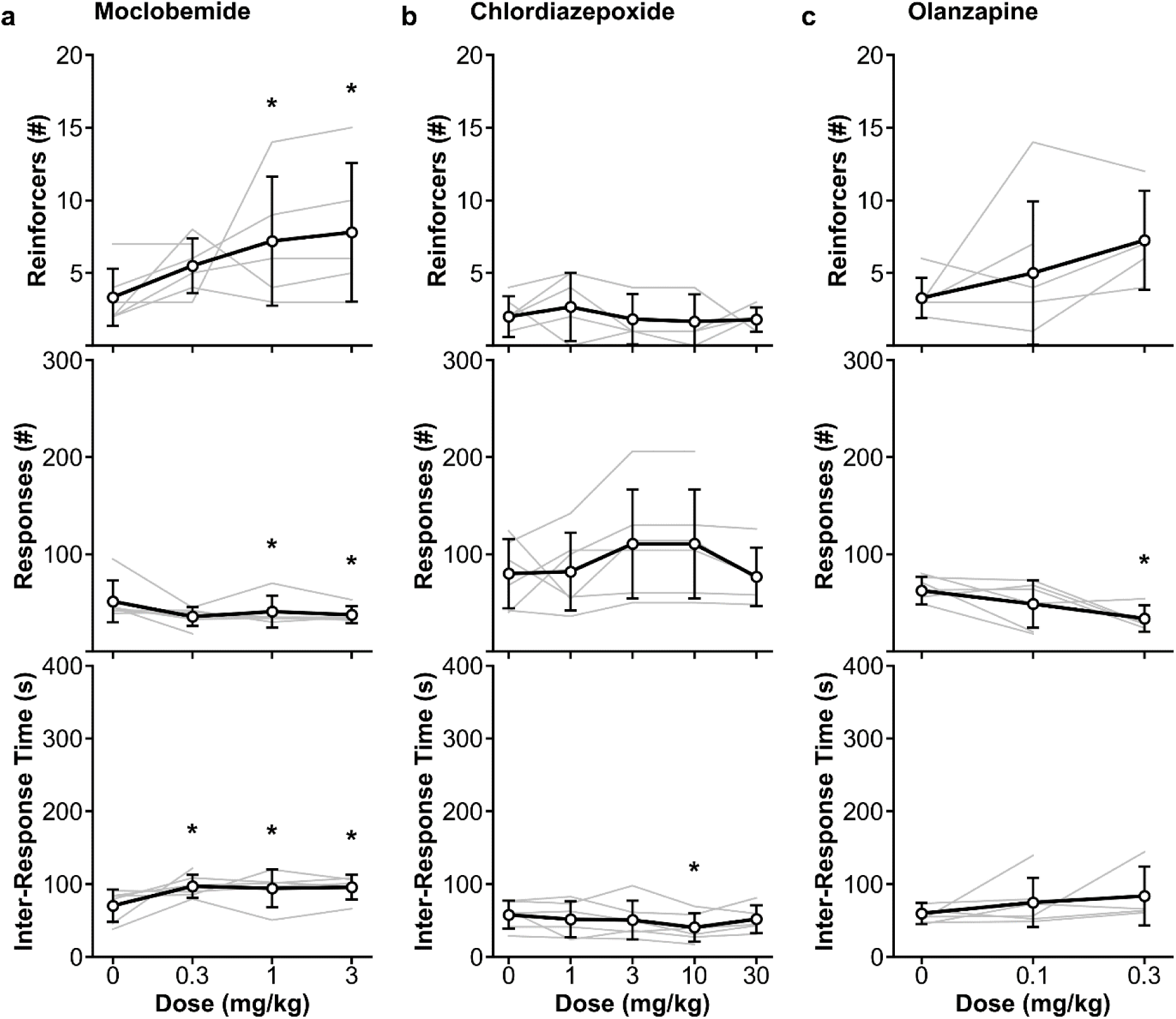
Behavioral effects of an MAOI and control compounds in the primate DRL task. Reinforcers earned, number of responses, and inter-response time following acute administration of (a) moclobemide, (b) chlordiazepoxide, and (c) olanzapine. The MAOI antidepressant moclobemide produced an antidepressant-like profile, whereas chlordiazepoxide showed no significant main effects. Olanzapine altered number of responses without producing a full antidepressant-like profile. Black lines are group means ± SD. Grey lines are individual animals. *p < 0.05 vs vehicle.

### 3.2. Pharmacological Specificity of the NHP-DRL Assay

To test the pharmacological specificity of the DRL task, we evaluated a set of compounds not primarily classified as antidepressants, including a benzodiazepine, an atypical antipsychotic, and psychostimulants. With a few exceptions, the resulting behavioral profiles were inconsistent with a full antidepressant-like pattern of DRL performance.

The benzodiazepine chlordiazepoxide (1–30 mg/kg; n = 6) did not significantly affect any of the dependent variables measured (Figure 3b; RF: F(4, 18.9) = 0.54, p = 0.708; RR: (F(4, 19) = 1.68, p = 0.195; IRT: F(4, 19) = 2.37, p = 0.089).

The atypical antipsychotic olanzapine (0.1–0.3 mg/kg; n = 7) produced a significant main effect on RR (Figure 3c; F(2, 7.4) = 4.79, p = 0.046), but not on RF (F(2, 9.0) = 3.57, p = 0.072) or IRT (F(2, 7.2) = 1.66, p = 0.256). The main effect of olanzapine on RR was driven by a significant decrease in performance with the 0.3 mg/kg dose compared to vehicle (p < 0.05).

To evaluate how stimulant compounds influenced DRL performance, we tested four psychostimulants with diverse mechanisms of action: nicotine, cocaine, methylphenidate, and modafinil.

Nicotine (0.03–0.3 mg/kg; n = 7) significantly affected all behavioral outcomes, with main effects on RF (Figure 4a; F(4, 22.8) = 3.22, p = 0.031), RR (F(4, 23) = 3.54, p = 0.022), and IRT (F(4, 23) = 3.46, p = 0.024). Post-hoc analyses revealed that these main effects were driven by the 0.3 mg/kg dose of nicotine which significantly increased RF and IRT, and decreased RR compared to vehicle treatment (all p < 0.05). Overall, nicotine shifted behavior in the same direction as classical antidepressants. However, because nicotine is not an antidepressant and stimulant compounds can alter DRL performance through multiple mechanisms, these findings warrant caution in interpreting this behavioral profile as uniquely indicative of antidepressant activity.

**Figure 4.**
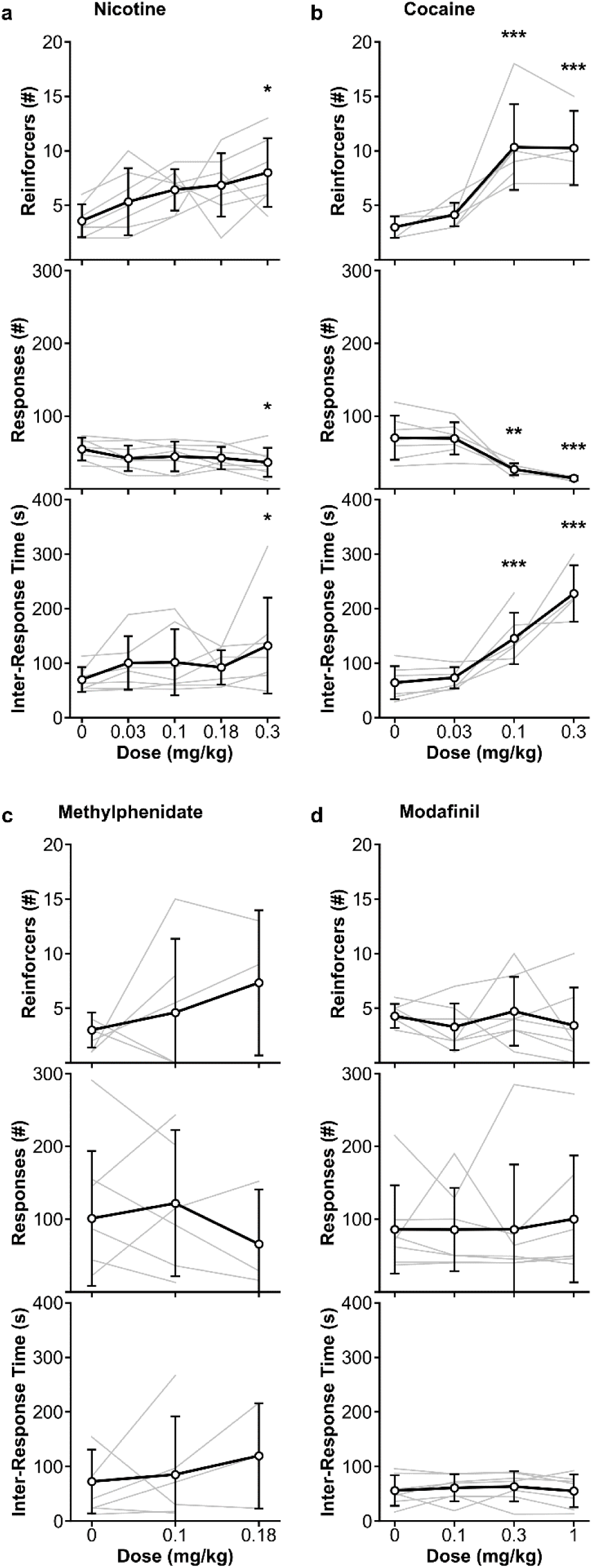
Effects of psychostimulant compounds on DRL task performance. Reinforcers earned, number of responses, and inter-response time following acute administration of (a) nicotine, (b) cocaine, (c) methylphenidate, and (d) modafinil. Nicotine and cocaine shifted DRL performance in an antidepressant-like direction, with increased reinforcers and inter-response time and reduced number of responses, whereas methylphenidate and modafinil produced no significant effects. Black lines are group means ± SD Grey lines are individual animals. *p < 0.05 vs vehicle. **p < 0.01 vs vehicle. ***p < 0.001 vs vehicle.

Cocaine (0.03–0.3 mg/kg; n = 7) produced robust, dose-dependent effects across all behavioral measures, with significant main effects on RF (Figure 4b; F(3, 14.2) = 44.43, p < 0.001), RR (F(3, 14.5) = 20.24, p < 0.001), and IRT (F(3, 14.2) = 24.57, p < 0.001). Post-hoc comparisons revealed the 0.1 and 0.3 mg/kg doses significantly increased RF, decreased RR, and increased IRT compared to vehicle administration (all p < 0.05). As with nicotine, this overlapping profile with antidepressants complicates a simple specificity interpretation and suggests that broad monoaminergic stimulation may be sufficient to improve DRL performance under the present task conditions.

In contrast to nicotine and cocaine, which shifted DRL performance in the same direction as antidepressants on several measures, the stimulants methylphenidate and modafinil did not significantly improve DRL performance. Methylphenidate (0.1–0.18 mg/kg; n = 8) did not significantly alter performance on any DRL measure. There were no significant main effects of dose on RF (Figure 4c; F(2, 11) = 0.39, p = 0.684), RR (F(2, 8.3) = 0.34, p = 0.724), or IRT (F(2, 8) = 0.34, p = 0.719).

Similarly, modafinil (0.1–1 mg/kg; n = 7) failed to produce significant effects on any behavioral measure in the DRL task. There were no significant main effects of dose on RF (Figure 4d; F(3, 18) = 1.21, p = 0.335), RR (F(3, 18) = 0.56, p = 0.645), or IRT (F(3, 18) = 0.48, p = 0.699).

### 3.3. Evaluation of PDE4 Inhibitors

Given growing interest in PDE4 inhibitors as potential antidepressants and their challenging side effect profile in clinical development, we tested three compounds with distinct selectivity and tolerability profiles: roflumilast, rolipram, and cilomilast. Each compound produced measurable effects on DRL performance, with emesis observed at higher doses across all agents.

Roflumilast (0.1–1 mg/kg; n = 7) produced significant main effects on all three behavioral measures, with increases in RF (Figure 5a; F(3, 15.4) = 4.19, p = 0.024), and IRT (F(3, 15.3) = 7.27, p = 0.003), and decreases in RR (F(3, 15.5) = 4.36, p = 0.021). Post-hoc analyses revealed significant behavioral changes at 0.3 mg/kg and 1 mg/kg. Specifically, IRT was significantly increased at both of these doses, RF was significantly increased at the 1.0 mg/kg dose, and RR was significantly decreased at the 1.0 mg/kg dose (all p < 0.05). Emesis was observed in six of the seven animals at the 1.0 mg/kg dose.

**Figure 5.**
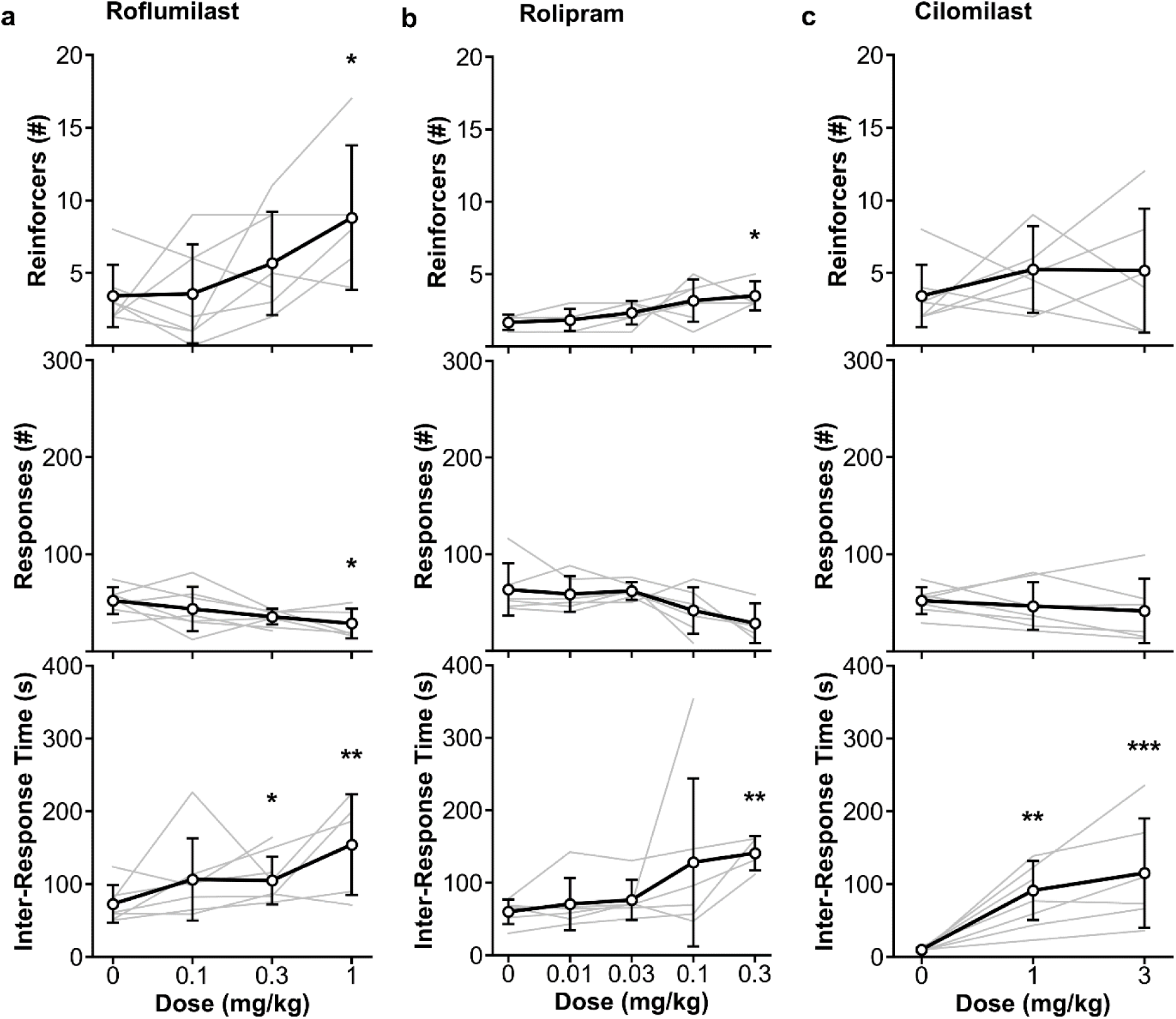
Effects of PDE4 inhibitors on DRL task performance. Reinforcers earned, number of responses, and inter-response time following acute administration of (a) roflumilast, (b) rolipram and (c) cilomilast. Roflumilast and rolipram produced antidepressant-like behavioral profiles, whereas cilomilast produced a partial behavioral effect limited to increased inter-response time. Black lines are group means ± SD. Grey lines are individual animals. *p < 0.05 vs vehicle. **p < 0.01 vs vehicle. ***p < 0.001 vs vehicle.

Rolipram (0.01–0.3 mg/kg; n = 6) significantly increased RF (Figure 5b; F(4, 23) = 3.69, p = 0.018) and IRT (F(4, 18.3) = 5.22, p = 0.006), but did not significantly affect RR (F(4, 18.2) = 2.60, p = 0.071). Post-hoc comparisons revealed that rolipram significantly increased RF and IRT at the 0.3 mg/kg dose (all p < 0.05). Notably, the 0.1 and 0.3 mg/kg doses were associated with emesis in all subjects.

Cilomilast (1–3 mg/kg; n = 7) administration had a significant main effect on IRT (Figure 5c; F(2, 9.9) = 19.74, p < 0.001), with significant increases following the 1 and 3 mg/kg doses compared to vehicle (all p < 0.05). Cilomilast did not significantly alter RF (F(2, 14) = 0.56, p = 0.586) or RR (F(2, 8.6) = 1.97, p = 0.198). All subjects exhibited emesis at the 3 mg/kg dose of cilomilast.

## 4. DISCUSSION

In this study, we adapted the DRL paradigm for use in NHPs and systematically evaluated its predictive validity, pharmacological sensitivity, and translational utility for detecting antidepressant-like effects. Antidepressants from multiple mechanistic classes elicited significant and consistent improvements in DRL performance. These effects were not consistently replicated by non-antidepressant compounds; however, nicotine and cocaine produced substantial overlap with the antidepressant-like behavioral profile, indicating that the model is highly sensitive to pharmacological modulation of DRL performance but not absolutely specific for antidepressants as a therapeutic class. Critically, we extended the application of this model to evaluate PDE4 inhibitors, a mechanistically novel antidepressant class whose clinical development has been limited by a narrow therapeutic window and emetic liability. The NHP model allowed us to assess both behavioral efficacy and emetic liability in the same subjects, offering a translational bridge between preclinical screening and clinical development. Together, these findings support the NHP DRL paradigm as a translational platform for preclinical evaluation of novel antidepressant therapies, particularly when both behavioral efficacy and tolerability are of interest.

### 4.1. Cognitive Processes Underlying DRL Performance

In the DRL task, subjects are reinforced for responding only after withholding behavior across an un-signaled delay interval, making the task sensitive to both impulsivity and timing precision. All animals were first trained to a stable baseline, which, under vehicle treatment, was marked by moderate response rates, short inter-response times, and few reinforcers earned, reflecting impulsive responding and imprecise timing. Improvement from this baseline requires two key processes: (1) enhanced inhibitory control to suppress premature responses, and (2) improved internal time estimation to better align responses with the delay requirement. Optimal DRL performance, therefore, is defined by reduced responding, longer inter-response times, and a high number of reinforcers earned, indicating infrequent, well-timed responding. Most antidepressants produced simultaneous improvements across all three behavioral measures, suggesting coordinated engagement of both inhibitory control and timing. This pattern provides a clear and interpretable behavioral signature of antidepressant-like behavioral effects and reinforces the utility of the DRL task as a translational model for probing core cognitive processes disrupted in depression.

### 4.2. Interpreting Antidepressant-Like Effects in the DRL Model

#### 4.2.1. Selective Serotonin Reuptake Inhibitors (SSRIs)

Consistent with prior rodent studies, the NHP DRL model demonstrated strong sensitivity to SSRIs. All SSRIs (paroxetine, citalopram, and fluoxetine) produced dose-dependent increases in RF and IRT and decreased RR. These results closely align with prior studies in rodent DRL models, which have repeatedly demonstrated that SSRIs elicit antidepressant-like behavioral changes (Danysz et al., 1988; van Hest et al., 1992; Olivier et al., 1993; Seiden et al., 1985; Marek et al., 1989; Sokolowski and Seiden, 1999; Cousins and Seiden, 2000; Cousins et al., 2000; Richards et al., 1993a; Dekeyne et al., 2002). The consistency of these effects across species reinforces the primate DRL task’s validity as a translational tool that captures antidepressant efficacy.

#### 4.2.2. Serotonin-norepinephrine reuptake inhibitors (SNRIs)

Although SNRIs are widely prescribed for depression, the rodent DRL literature has largely failed to demonstrate antidepressant-like effects with this class. For example, venlafaxine failed to produce antidepressant-like effects in rodent DRL, even at doses that produce clear effects in other antidepressant-sensitive assays (Dekeyne et al., 2002). In contrast, our findings demonstrate that the primate DRL model is sensitive to SNRI treatment. Both duloxetine and sibutramine significantly improved performance, increasing RF and IRT while decreasing RR.

These species differences may reflect broader pharmacological or neurobiological divergences in monoaminergic regulation of response inhibition and interval timing. They may also highlight distinctions between individual drugs within the SNRI class. For instance, duloxetine has a 10-fold selectivity for serotonin over norepinephrine, while venlafaxine exhibits a 30-fold preference (Montgomery, 2008), potentially explaining duloxetine’s more balanced noradrenergic contribution to DRL performance. Clinically, duloxetine has been shown to outperform SSRIs in some populations, particularly individuals with treatment-resistant depression (Mallinckrodt et al., 2007; Pitchot et al., 2010; Rodrigues-Amorim et al., 2020). However, other findings indicate that duloxetine may not be as effective as venlafaxine (Schueler et al., 2011).

Sibutramine, while structurally and mechanistically similar to antidepressants, was originally developed as an appetite suppressant and is not approved for depression (Araujo & Martel, 2012; Bray et al., 1999). Nonetheless, its dual blockade of serotonin and norepinephrine reuptake mirrors that of traditional SNRIs (Stahl et al., 2014). Our results show that sibutramine improves DRL performance, suggesting that its underlying pharmacology is sufficient to engage the same cognitive processes targeted by approved antidepressants. This supports the utility of the primate DRL model in detecting compounds with dual reuptake inhibition, even when their primary clinical indication lies outside of depression.

#### 4.2.3. Selective norepinephrine reuptake inhibitors (NRIs)

NRIs offer an important test of the DRL model’s sensitivity to noradrenergic modulation independent of serotonergic mechanisms. We evaluated two NRIs, reboxetine and atomoxetine, that differ in clinical application and efficacy (Kobayashi et al., 2010; Holm & Spencer, 2012; Ding et al., 2014). Reboxetine produced dose-dependent improvements on DRL performance across all behavioral endpoints. These results are consistent with previous literature utilizing this compound in the rodent DRL model (Dekeyne et al., 2002; Wong et al., 2000), but contrast with the questionable efficacy of reboxetine as an antidepressant in humans (Cipriani et al., 2009). In contrast, atomoxetine had modest effects on DRL performance, significantly increasing RF at the mid and high doses, while no effects emerged for IRT or RR. These findings align with the fact that atomoxetine is approved for clinical use in attention-deficit/hyperactivity disorder, not depression (The Atomoxetine ADHD and Comorbid MDD Study Group et al., 2007; Michelson et al., 2003). It is likely that its relatively weak affinity for the norepinephrine transporter compared to other NRIs explains the lack of robust behavioral modulation across the dependent variables (Aldosary et al., 2022). The divergence between reboxetine and atomoxetine emphasizes that not all NRIs exert equivalent effects in the DRL task. These results also suggest that norepinephrine reuptake inhibition can modulate DRL behavior, but only under specific neurochemical or receptor engagement conditions.

#### 4.2.4. Norepinephrine-dopamine reuptake inhibitor (NDRI)

The NDRI bupropion significantly improved DRL performance by increasing IRT and reducing RR, though the increase in reinforcers earned did not reach statistical significance. These effects were most pronounced at the highest dose tested; however, the magnitude of these effects was relatively mild compared to other antidepressants. This parallels the rodent literature where bupropion has produced weak and inconsistent effects on DRL tasks (Dekeyne et al., 2002; Seiden et al., 1985; Zhang et al., 2024). This relatively limited efficacy may reflect bupropion’s pharmacological profile, which includes weaker inhibition of norepinephrine and dopamine transporters compared to classic NRIs or stimulants, and an absence of serotonergic activity (Stahl et al., 2004). While clinically effective for some individuals, especially those with atypical or treatment-resistant depression, bupropion is also often used as an adjunctive therapy, suggesting that its modest behavioral impact in the DRL task may be pharmacologically appropriate (Raju et al., 2025; Nasr et al., 2014).

#### 4.2.5. Tricyclic antidepressant (TCA)

Tricyclic antidepressants are among the earliest pharmacologic treatments for depression (Kamp et al., 2024). Although their clinical use has declined due to unfavorable side effect profiles, their mechanisms, primarily inhibition of norepinephrine reuptake, remain relevant for evaluating antidepressant efficacy in preclinical models. In the present study, the TCA desipramine significantly increased RF but did not change RR or IRT, supporting a partial antidepressant-like behavioral profile.

These findings are consistent with rodent literature demonstrating desipramine-induced improvements in DRL performance (O’Donnell and Seiden, 1983; Danysz et al., 1988; Britton and Koob, 1989; Dekeyne et al., 2002; Richards and Seiden, 1991; Richards et al., 1993b; O’Donnell and Seiden, 1984; Jackson et al., 1995; Richards et al., 1993a). There are, however, some differences between our results and those from rodents. In rats, desipramine typically produces robust increases in RF and IRT along with decreases in RR. While our primate data mirrored the increased RF, we saw no effect of desipramine on RR or IRT. Additionally, whereas primates showed an increase in RF following desipramine treatment, one rodent study found no effect of desipramine on RF (Jackson et al., 1995). Finally, whereas in primates the increase in RF was driven by the highest dose tested, rodent studies have frequently reported broader dose responsiveness. These discrepancies may reflect species-specific sensitivity or suggest that higher doses are required to elicit full effects in macaques. Overall, while broadly concordant with rodent data, the primate DRL model may provide a more conservative or selective measure of TCA efficacy.

#### 4.2.6. Monoamine Oxidase Inhibitor (MAOI)

MAOIs were among the first antidepressants developed but are less commonly prescribed today due to safety concerns related to medication and dietary tyramine interactions (Brown et al., 1989; Edinoff et al., 2022). Nonetheless, MAOIs remain mechanistically informative and are particularly useful for treatment-resistant depression (Yamada and Yasuhara, 2004). In the present study, the reversible MAOI moclobemide produced a robust antidepressant-like profile in the primate DRL model. These effects were dose-dependent and consistent across all three measures. Importantly, these findings align with rodent DRL studies using irreversible MAOIs such as phenelzine, iproniazid, and isocarboxazid (O’Donnell and Seiden, 1982; 1983). However, our results extend this literature by demonstrating that a reversible, MAO-A-selective inhibitor can produce similar behavioral improvements. This is clinically relevant because reversible MAOIs like moclobemide have a more favorable safety profile than irreversible agents, yet still may preserve the therapeutic benefit seen in classical MAOI compounds (Nair et al., 1993). The efficacy of moclobemide in this primate model suggests that enhancing monoaminergic tone through selective MAO-A inhibition is sufficient to produce antidepressant-like effects, and further supports the use of primate DRL as a translational tool to evaluate mechanistically diverse antidepressants.

#### 4.2.7. Benzodiazepines (BZD) and Antipsychotics

To assess the pharmacological specificity of the DRL paradigm, we included compounds not traditionally considered antidepressants. The benzodiazepine chlordiazepoxide failed to improve DRL performance, producing no significant changes in RF, RR, or IRT. This aligns with prior rodent DRL studies demonstrating that benzodiazepines often either impair performance or increase impulsivity, likely due to their sedative and disinhibitory effects (van Hest et al., 1992; Richards et al., 1993b; O’Donnell and Seiden, 1982).

The atypical antipsychotic olanzapine elicited a significant main effect only on RR, a behavioral pattern not matching the profile typically observed with antidepressants. Olanzapine has yet to be tested in the rodent DRL model, but clozapine, which is structurally and pharmacologically similar, has been tested a handful of times. Specifically, clozapine decreased RR and RF (Seiden et al., 1985), which is partially similar to the primate olanzapine results reported here. Similarly, acute clozapine failed to rescue deficits in DRL produced by administration of phencyclidine (Compton et al., 2001).

#### 4.2.8. Psychostimulants

We tested four stimulant compounds with distinct pharmacological mechanisms to determine whether behavioral activation alone is sufficient to improve DRL performance, or whether antidepressant-like effects depend on specific neuromodulatory engagement (Pary et al., 2015; Hai et al., 2022). Nicotine and cocaine both produced robust antidepressant-like profiles on the task. Although their primary mechanisms differ, with nicotine acting through nicotinic acetylcholine receptors and cocaine blocking monoamine reuptake transporters, both increase extracellular dopamine, norepinephrine, and serotonin (Ruiz et al., 2021; Beer, 2016; Wignall & de Wit, 2011). This shared profile of broad monoaminergic activation may account for their convergent behavioral effects. Importantly, these findings indicate that improved DRL performance in this primate preparation is not uniquely restricted to clinically approved antidepressants. Rather, under the present task conditions, broad monoaminergic enhancement may be sufficient to improve the inhibitory control and temporal regulation required for optimal DRL performance.

In contrast, methylphenidate and modafinil failed to improve DRL performance. Both compounds lack meaningful serotonergic activity (Repantis et al., 2010), which may limit their ability to engage the behavioral processes required for antidepressant-like effects on the task. Methylphenidate primarily blocks dopamine and norepinephrine reuptake, but its dopaminergic predominance may increase arousal and motor output without improving response regulation (Schrantee et al., 2016). This interpretation is consistent with rodent findings that methylphenidate can increase impulsivity and disrupt performance on tasks requiring response inhibition (Navarra et al., 2008; Richards et al., 1993b). Modafinil promotes wakefulness through orexin and dopamine systems, but has minimal impact on serotonin or norepinephrine and may therefore fail to engage the mechanisms most relevant for improved DRL performance (Gerrard & Malcolm, 2007; Mereu et al., 2014). Together, these findings suggest that the primate DRL model is highly sensitive to antidepressant-relevant behavioral change, although its pharmacological specificity is not absolute and depends on both mechanism of action and task design.

### 4.3. Effects of Phosphodiesterase-4 (PDE4) Inhibitors

Having validated the primate DRL model as sensitive to compounds with antidepressant-relevant monoaminergic activity, we next applied it to a novel drug class. Phosphodiesterase-4 inhibitors such as roflumilast, rolipram, and cilomilast have shown promising antidepressant properties in preclinical studies (Wang et al., 2015, 2020; Yu et al., 2021). However, a major limitation in the development of clinical PDE4 compounds is dose-limiting side effects. We used the primate DRL model to evaluate both the antidepressant-like efficacy and tolerability of these compounds in a translationally relevant context.

Importantly, NHPs offer a distinct advantage over rodents for evaluating PDE4 inhibitors. While rodents are widely used for early antidepressant screening, they lack a gag reflex and cannot vomit, limiting their utility in detecting dose-limiting side effects such as emesis, a major limitation in the development of clinical PDE4 compounds. The primate DRL model thus offers a unique opportunity to simultaneously assess efficacy and tolerability in a single behavioral paradigm, improving predictive validity for human clinical trials.

Two of the compounds, roflumilast and rolipram, exhibited classic antidepressant-like profiles in the DRL task. These effects mirror prior rodent studies showing that PDE4 inhibitors improve DRL performance (Zhang et al., 2006; O’Donnell, 1993; O’Donnell and Frith, 1999; Zhang et al., 2005; O’Donnell and Zhang, 2004). In contrast, cilomilast only increased IRT, and failed to affect RF or RR, yielding an incomplete antidepressant-like profile.

Despite these drug efficacy signals, tolerability remains a major limitation. Emesis was observed in all monkeys administered rolipram at 0.1 or 0.3 mg/kg, in all subjects given 3 mg/kg cilomilast, and in six of seven animals treated with 1 mg/kg roflumilast. These side effects closely resemble clinical experiences with PDE4 inhibitors, where nausea and vomiting curtailed further development despite clear antidepressant potential (Zeller et al., 1984; Burnouf et al., 1998). Thus, while the DRL model detects antidepressant efficacy for PDE4 inhibitors, it also recapitulates their dose-limiting side effect profile, emphasizing the need for improved tolerability in this drug class.

### 4.4. Drug Repurposing

An additional strength of the NHP DRL model is its potential utility for both drug repurposing and novel therapeutic discovery. Several non-traditional compounds tested here produced behavioral effects overlapping with antidepressant-like behavioral profiles, illustrating how this assay may help identify antidepressant-relevant properties in approved or previously developed agents. At the same time, the model was sensitive to mechanistically novel PDE4 inhibitors and, importantly, detected dose-limiting emesis in the same subjects. This combination of efficacy-related and tolerability-related readouts may be particularly valuable when prioritizing compounds for further development.

### 4.5. Limitations

Several features of the present design should be considered when interpreting pharmacological specificity. First, unlike standard rodent DRL procedures that typically use a fixed interval across subjects, the DRL delay was individualized in the present NHP studies to maintain stable within-subject performance over long testing periods and to minimize floor effects. Although this approach increased sensitivity to repeat pharmacological testing within subjects, it reduces direct comparability with fixed-interval rodent procedures and may have influenced the behavioral expression of stimulant compounds. Second, baseline performance was intentionally maintained in a relatively narrow range of 2–6 reinforcers per session, which may have limited our ability to detect the more classically disruptive stimulant profile often observed in rodent DRL studies. These features may help explain why nicotine and cocaine shifted performance in the same direction as antidepressants in the present study. Accordingly, we interpret the present findings as evidence of strong translational sensitivity to antidepressant-relevant pharmacology, while recognizing that pharmacological specificity in this NHP preparation is shaped by schedule design and baseline conditions.

### 4.6. Conclusion

The present findings support the DRL task as a translationally relevant model for evaluating antidepressant-like effects in non-human primates. Across multiple antidepressant classes, including SSRIs, SNRIs, NRIs, NDRIs, TCAs, MAOIs, and PDE4 inhibitors, the task detected a convergent behavioral pattern characterized by increased reinforcers earned, lengthened inter-response times, and reduced responding. Although this pattern was not consistently observed across non-antidepressant drug classes, nicotine and cocaine produced substantial overlap with the antidepressant-like profile, indicating the model is highly sensitive to pharmacological modulation of DRL performance but not absolutely specific for antidepressants as a therapeutic class. Importantly, the DRL model also captured clinically relevant tolerability signals, including emesis with PDE4 inhibitors, an outcome not detectable in rodent models. By enabling simultaneous evaluation of behavioral efficacy and side effect liability, this model provides a valuable framework for translational antidepressant screening and preclinical drug development.

## Data Availability Statement

The datasets generated during the current study are available from the corresponding author on reasonable request.

## Acknowledgments

We are grateful to Joseph Cefalu and Hung Dao for their assistance with data collection. This work was supported by F. Hoffmann-La Roche.

## Author Contributions

JAV, JLB, and JGW contributed to the conception and design of the work, JLB, JAV and CGB collected data, CRV, SRD, and CGB analyzed the data; CRV, SRD, and CGB interpreted the data and drafted the manuscript; JAV and JGW reviewed and edited the manuscript, all authors approved the final version for publication.

